# Coordination between discrete Mitotic Arrest Deficient 1 (MAD1) domains is required for efficient mitotic checkpoint signaling

**DOI:** 10.1101/212704

**Authors:** Wenbin Ji, Yibo Luo, Ejaz Ahmad, Song-Tao Liu

**Affiliations:** From the Department of Biological Sciences, University of Toledo, Toledo, OH 43606

**Keywords:** mitosis, signal transduction, kinetochore, mitotic spindle, mitotic checkpoint, checkpoint control, MAD1, MAD2, MPS1 kinase

## Abstract

As a sensitive signaling system, the mitotic checkpoint ensures faithful chromosome segregation by delaying anaphase onset when even a single kinetochore is unattached. The key signal amplification reaction for the checkpoint is the conformational conversion of open MAD2 (O-MAD2) into closed MAD2 (C-MAD2). The reaction was suggested to be catalyzed by an unusual catalyst, a MAD1:C-MAD2 tetramer, but how the catalysis is executed and regulated remains elusive. Here we report that in addition to the well-characterized middle region (MIM), both amino- and carboxyl-terminal domains (NTD and CTD) of MAD1 also contribute to the mitotic checkpoint. In contrast to MIM that stably associates with C-MAD2, MAD1-NTD and CTD surprisingly bind to both O-MAD2 and C-MAD2, suggesting their interactions with substrates and products of the O-C conversion. MAD1-NTD also interacts with CTD. MPS1 kinase interacts with and phosphorylates both NTD and CTD. The phosphorylation reduces the NTD:CTD interaction and CTD interaction with MPS1. Mutating CTD phosphorylation sites including Thr716 compromises MAD2 binding and the checkpoint responses. Ser610 and Tyr634 also contribute to the checkpoint. Our results have uncovered previously unknown interactions of MAD1-NTD and CTD with MAD2 conformers and their regulation by MPS1 kinase, providing novel insights into the mitotic checkpoint signaling.

---

The mitotic checkpoint is a crucial signal transduction pathway that contributes to faithful chromosome segregation (1–4). A single unattached kinetochore delays anaphase onset, underscoring the importance of signal amplification for the mitotic checkpoint (5). The conversion of MAD2 from open (O-MAD2) to closed (C-MAD2) conformation is a well-recognized signal amplification mechanism for the mitotic checkpoint (6,7). O-MAD2 is the predominant conformer in interphase cells (8,9). During prometaphase intracellular C-MAD2 concentration is increased to promote formation of the mitotic checkpoint complex (MCC), which binds and inhibits the anaphase promoting complex/cyclosome (APC/C) (1–4). In the current model, the MAD2 O-C conversion is catalyzed by a 2:2 MAD1:C-MAD2 tetramer localized at unattached kinetochores, whereby cytoplasmic O-MAD2 hetero-dimerizes with the C-MAD2 moiety in the catalyst and morphs into C-MAD2 through unknown mechanism but possibly involving some intermediate folding states (I-MAD2) (6,7,10).

Two major questions remain unanswered for the model. First, human MAD1 is a mitotic checkpoint protein of 718 amino acid residues but the formation of a 2:2 heterotetramer with C-MAD2 involves only its MAD2 interaction motif (MIM, 485-584 residues) especially a disordered loop spanning 530-550 residues (Fig. 1a) (11). However, some earlier and recent experiments argued for functional importance of the C-terminal domain of MAD1 (585-718 residues, MAD1^CTD^) in maintaining the mitotic checkpoint (12–18).

**Fig. 1.**
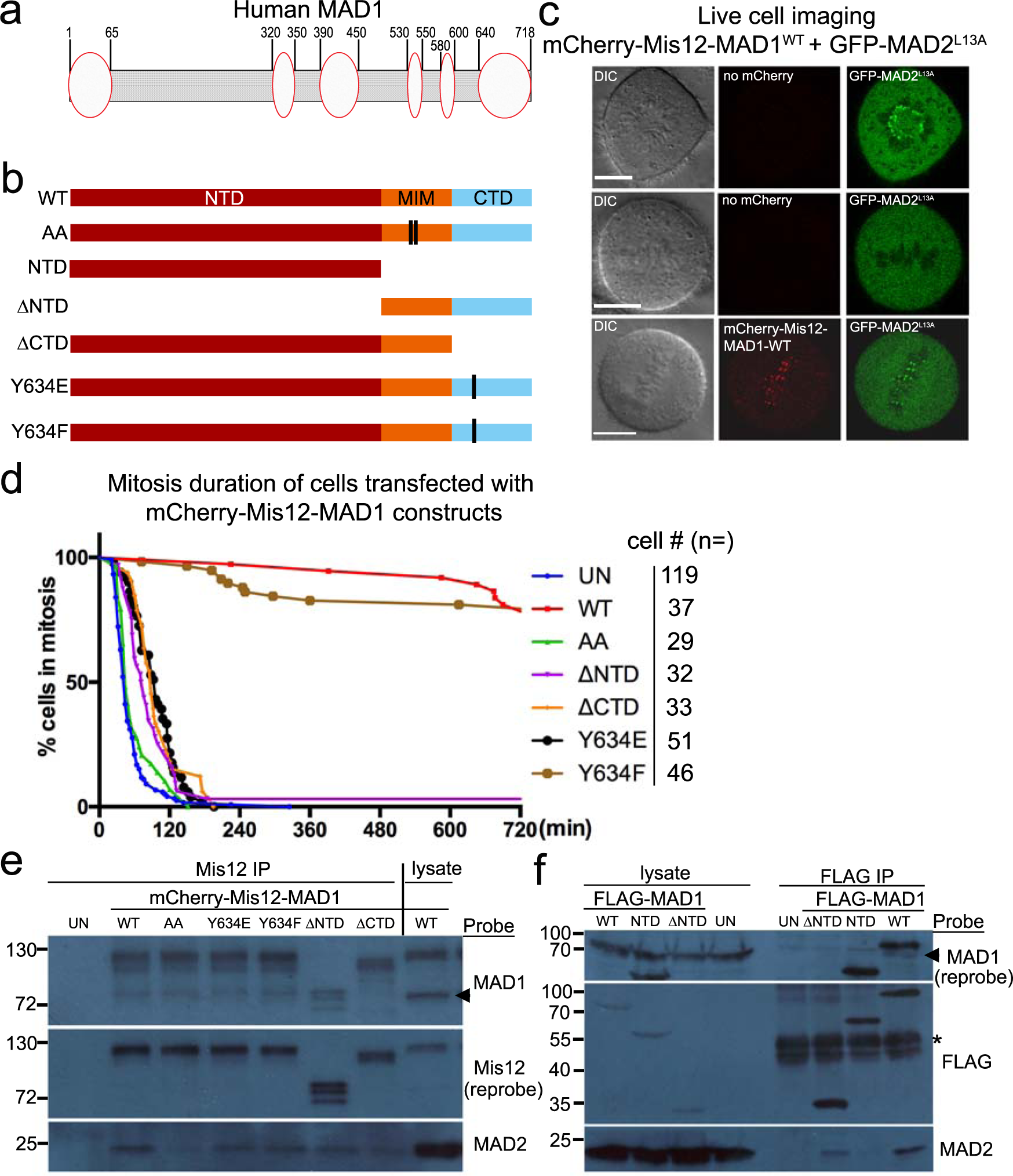
Both MAD1-NTD and CTD are required for MAD1 activity. **(a)** Diagram of human MAD1 showing that multiple segments of MAD1 may form structures (ovals) other than coiled coil (shaded area). **(b)** Shown are various MAD1 mutants and truncations used for fusion with mCherry-Mis12. NTD: N-terminal domain; MIM: MAD2 interacting motif; CTD: C-terminal domain. **(c)** Images of live cells either transfected with GFP-MAD2^L13A^ alone or together with mCherry-Mis12-wild type (WT) MAD1. Bar=10 µm. **(d)** Mitotic durations of HeLa cells either untransfected (un) or transfected with mCherry-Mis12 fused with MAD1-WT, AA (defective in MAD2 binding due to mutations in MIM), ∆NTD, ∆CTD, Y634E, or Y634F were shown. Cell numbers imaged for each construct were listed on the right. **(e)** Lysates from control (UN) or transfected cells were subjected to anti-Mis12 immunoprecipitation followed by Western blots for Mis12-MAD1 fusions (probed with anti-Mis12 antibody), MAD2 and endogenous MAD1. Molecular weight markers are marked on the left (in kD). The arrow indicates the position of endogenous MAD1. **(f)** Lysates from control (UN) or FLAG-MAD1 transfected cells were subjected to anti-FLAG immunoprecipitation to probe for MAD1 and MAD2. The asterisk indicates IgG heave chain.

Moreover, even the MAD1 fragment encompassing (485-718) residues only exhibits low catalytic activity for MAD2 O-C conversion *in vitro* (12,13,19). The N-terminal domain of MAD1 (1-485 residues, MAD1^NTD^) was thought to target the protein to nuclear envelope or kinetochores but may also interact with other proteins (20–23). Whether and how MAD1 domains outside MIM contribute to the checkpoint signaling warrants re-visit and careful investigation. Second, MAD1 forms a cell cycle independent complex with C-MAD2; how the complex only becomes an effective catalyst during prometaphase needs to be better defined (18,24). Several kinetochore-localized mitotic kinases including MPS1 kinase were known to elevate C-MAD2 production but direct biochemical evidence is still incomplete despite exciting recent progress (25–31).

Here we report our results targeting the above two questions. While our manuscript was prepared, two reports were published indicating MPS1 phosphorylating MAD1 to enhance MAD2 O-C conversion and the MCC assembly (29,30). Our results support the importance of MAD1 Thr716 phosphorylation, but we have also uncovered other protein-protein interactions among MAD1, MAD2 and MPS1 and their phosphorylation dependent regulation. Our work highlights the coordination of different MAD1 domains in efficient mitotic checkpoint signaling and provides further mechanistic insights into the O-C conversion reaction.

## RESULTS

### MAD1 N-terminal and C-terminal domains (NTD and CTD) are required for efficient mitotic checkpoint signaling

In studying the MAD2 O-C conversion, earlier work has detailed the conformational changes of MAD2 (6,32). We reasoned that better characterization of the MAD1:C-MAD2 catalyst would provide further mechanistic insights into the conversion reaction and hence the signal amplification step of the mitotic checkpoint. We noted that even though MAD1 is commonly depicted as a rigid coiled coil protein, parts of its NTD and CTD have been shown or were predicted to remain disordered or adopt other structures (Fig. 1a) (Fig. S1)(11,33,34). We first investigated possible contribution of MAD1^NTD^ and MAD1^CTD^ to the mitotic checkpoint using a “separation-of-function” system developed by Maldonado and Kapoor (26). In this system, a mCherry-Mis12-MAD1 fusion construct was exploited to examine “catalytic efficiency” of the MAD1:C-MAD2 catalyst without concerns over the “kinetochore targeting” aspect of its regulation (26) (Fig. 1b). Although endogenous MAD1 and MAD2 disappear from metaphase kinetochores which presumably are occupied by spindle microtubules, expression of wild type MAD1 (MAD1^WT^) fused with “constitutive” kinetochore protein Mis12 retained MAD1 at metaphase plate and recruited GFP-MAD2^L13A^ to these metaphase kinetochores (26,28) (Fig. 1c, Fig. S2). MAD2^L13A^ is a MAD2 mutant locked in C-conformation (13,32). The persistence of MAD1 and MAD2 at metaphase attached kinetochores was sufficient to trigger a >12 hr mitotic arrest in HeLa cells (26,28)(Fig. 1d). The arrest is dependent on C-MAD2 binding to MAD1, as cells expressing the fusion with MAD2-binding deficient MAD1^AA^ mutant (K541A, L543A in MIM) finished mitosis within ~60 min on average (Fig. 1d, Fig. S2) (26). Note no GFP-MAD2^L13A^ is localized at metaphase kinetochores containing mCherry-Mis12-MAD1^AA^, although GFP-MAD2^L13A^ does appear at the last few unattached kinetochores most likely due to presence of endogenous MAD1 there (Fig. S2, compare the second and third columns). Furthermore, co-expression of MAD2^ΔC10^, an O-conformer locked mutant of MAD2 (6,7), abolished the mitotic arrest in MAD1^WT^ transfected cells (data not shown), corroborating that the arrest is due to O-C conversion dependent checkpoint responses (28,35,36). In consistence with previous reports (14–17), MAD1 missing (597-718) residues (MAD1^ΔCTD^), even as a fusion with Mis12, could not maintain mitotic arrest (98±7 vs 749±22 min for MAD1^WT^, mean±SD, P<0.0001, student’s *t*-test, there might be an underestimation for MAD1^WT^ transfected cells as the movies last 13 hrs). Moreover, a specific MAD1^Y634E^ mutant also abolished the mitotic arrest, while a MAD1^Y634F^ mutant did not significantly impact mitotic duration (Fig. 1d). Y634 is situated close to the junction between the coiled coil subdomain (597-637 residues) and the globular subdomain (638-718 residues) of the MAD1^CTD^ (33). We noticed this site during screening potential MAD1 phospho-mutants as Y634 was reported to be phosphorylated *in vivo* (37). Interestingly, MAD1 missing (1-485) residues (MAD1^ΔNTD^) cannot maintain prolonged mitosis either (average duration: 101±22 min, Fig 1d).

When examined by Mis12 immunoprecipitation, in addition to MAD1^AA^ failing to recruit MAD2, slight reduction of steady levels of MAD2 binding to other MAD1 mutants were also noticed (Fig.1e). The reduction of MAD2 binding became more obvious for some mutants when FLAG immunoprecipitation was performed using cells transfected with FLAG-MAD1 constructs (Fig 1f). In addition, MAD1^ΔCTD^ and MAD1^ΔNTD^ cannot interact with endogenous MAD1, indicating a dimerization defect (Fig 1e&f, note that mCherry-Mis12-MAD1^ΔNTD^ runs at the same position as endogenous MAD1. More on MAD1 dimerization in the last section of RESULTS). Taken together, results shown in Figure 1 confirm the important role of MAD1^MIM^, but also reveal that both NTD and CTD of MAD1 are required for an efficient mitotic checkpoint.

### MAD1-NTD and -CTD bind to both O-MAD2 and C-MAD2

We hypothesized that the NTD and CTD of MAD1 facilitate mitotic checkpoint responses by enhancing MAD2 O-C conversion. Based on the analogy to an isomerase which at least transiently interacts with its substrate and product, we prepared recombinant proteins and examined potential interactions between GST-tagged MAD1-NTD or CTD with untagged O-or C-MAD2, supplied as MAD2^ΔC10^ or MAD2^L13A^ conformation-locked mutants respectively (6,7,36) (Fig. 2a). GST-pulldown results showed that NTD and CTD bind to both conformers of MAD2 (lanes 1, 3 and 5, 7 in Fig. 2b). Under similar conditions, the MAD1^MIM^ only binds to C-MAD2 just as previously reported (compare MAD2 in Fig 2b, lanes 2&6) (11,38). The conformational status of the O- and C-MAD2 mutants were further verified that only MAD2^L13A^ binds to GST-CDC20^(111-138)^ or GST-BUBR1^(1-371)^, also as reported before (36,38) (Fig. S3). The novel interactions were not mediated by tags, as GST alone did not pull down any MAD2 (Fig. 2b, lanes 4&8). In addition, GFP-MAD2^ΔC10^ was found to be recruited to centromeres in interphase cells expressing mCherry-Mis12-MAD1 fusions, supporting that the interaction between O-MAD2 with MAD1, although surprising, could happen in cells (Fig. 2c, MAD1^ΔCTD^ used here). Maintaining MAD1 fragments at the endogenous concentration of 60 nM, titrating GST-pulldown experiments estimated that half maximal binding concentrations (BC50) for NTD, MIM and CTD interactions with C-MAD2 and O-MAD2 all fall in the range of 100 ~ 250 nM (Fig. S3).

**Fig. 2.**
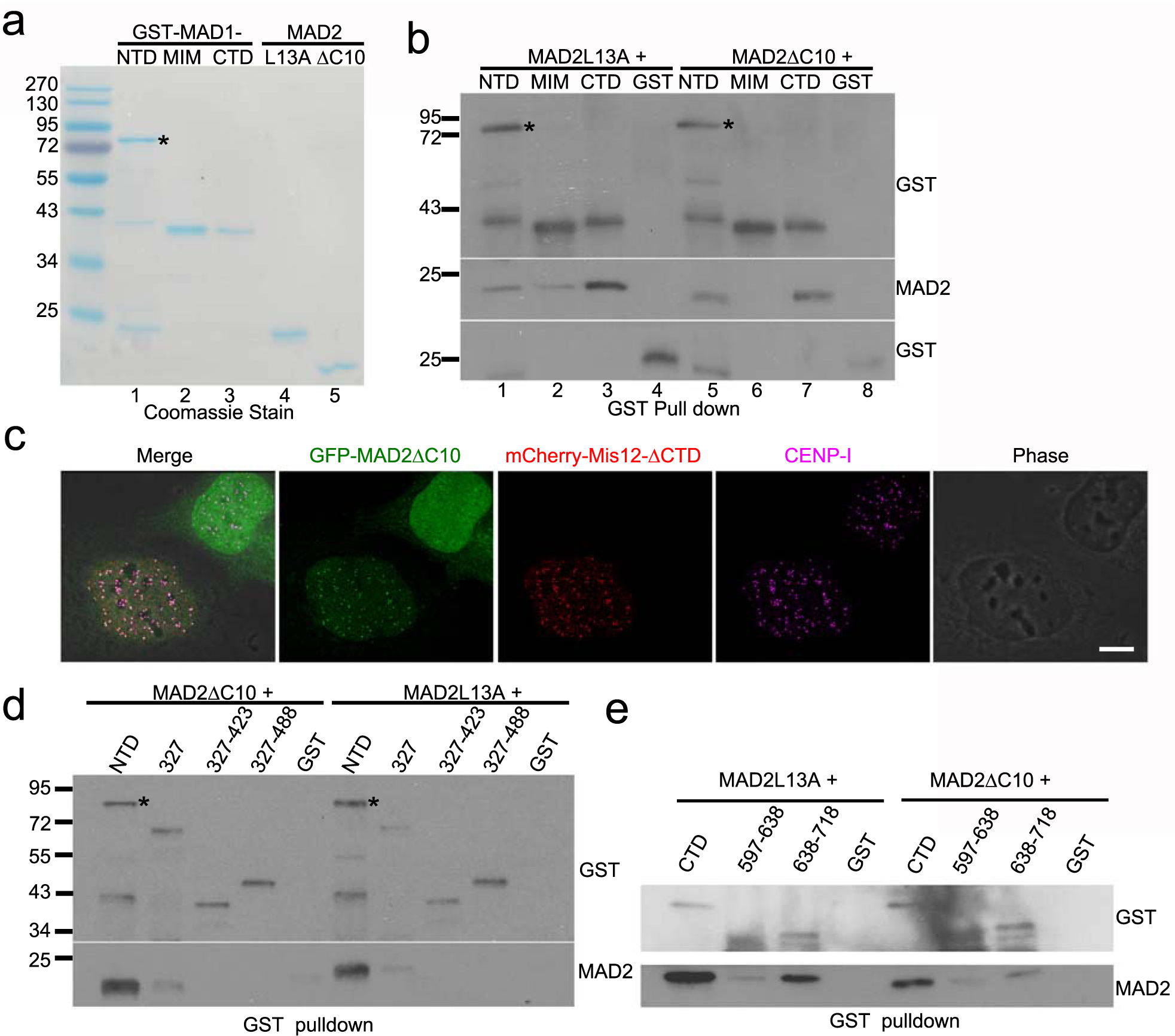
MAD1-NTD and CTD interact with both O-MAD2 and C-MAD2. **(a)** Coommassie stain after SDS-PAGE of purified recombinant proteins. GST-NTD tends to be more labile for degradation. The asterisk marks expected size of GST-NTD. **(b)** GST-tagged MAD1-NTD, MIM, CTD or GST alone was incubated with either MAD2^L13A^ or MAD2^∆C10^, and GST pull-down assays were performed followed by Western blot of GST and MAD2. **(c)** Immunofluorescence of two interphase cells transfected with GFP-MAD2^∆C10^. Note that the GFP signals are recruited to centromeres only in the cell co-expressing mCherry-Mis12-MAD1^ΔCTD^. Centromeres are stained with anti-CENP-I antibody. Bar=10 µm. **(d)** GST-tagged MAD1 N-terminal truncations were incubated with MAD2^∆C10^ or MAD2^L13A^ followed by GST pull-down assays. **(e)** GST-tagged MAD1-CTD coiled-coil subdomain (597-638 residues) or globular subdomain (638-718 residues) were incubated with MAD2^∆C10^ or MAD2^L13A^ followed by GST pull-down assays.

We attempted to further define the regions on MAD1-NTD or CTD responsible for association with MAD2. Several MAD1-NTD or CTD truncations produced either insoluble or heavily degraded proteins (data not shown). However, testing with MAD1-NTD truncations that we were able to purify, including MAD1^(1-327)^, MAD1^(327-423)^, and MAD1^(327-488)^, revealed dramatically reduced MAD2 binding capability of these fragments (Fig. 2d). The globular subdomain (638-718 residues) of MAD1-CTD retained approximately half MAD2 binding capacity of MAD1-CTD while the coiled coil subdomain in the CTD showed only residual binding (Fig. 2e). These combined results suggested that the integrity of MAD1-NTD or CTD is crucial for binding to O-MAD2 or C-MAD2.

### A novel interface in MAD2 is employed for its association with MAD1-NTD or –CTD

A MAD2 molecule has two well-characterized interfaces for protein-protein interactions: the “safety belt” characteristic of C-MAD2 conformation to which MAD1-MIM and CDC20 binds (11,38–40), and the dimerization domain (primarily αC helix) that allows MAD2 to form O:C or C:C dimers or interact with p31^comet^ or BUBR1 (13,32,36,41,42). MAD1-NTD and CTD bind to both MAD2^ΔC10^ and MAD2^L13A^, suggesting no discrimination against either MAD2 conformation (Fig. 2b). To further support this, wild type (WT) MAD2 existing as a mixture of both O and C conformers bind to all three MAD1 fragments, while another MAD2 mutant MAD2^S195D^, predominantly in O conformation (43), binds to MAD1-NTD or CTD but not MIM (Fig. 3a). On the other hand, MAD2^LARQ^ (L13A/R133E/Q134A), a dimerization defective C-MAD2 mutant, associates with MAD1-NTD or CTD at similar levels as MAD2^L13A^ (Fig. 3b). MAD2^ΔN10^, which might mimic an intermediate conformation during O to C-MAD2 conversion (10), binds to all three MAD1 fragments (Fig. 3c). The above results suggest that the safety belt (or C-conformation), the dimerization domain (at least the RQ mutant) or the N-terminal 10 amino acids are not essential for MAD2 to interact with MAD1-NTD or CTD. Therefore a novel interface on MAD2, possibly shared by both O- and C-conformers, is employed for interactions with MAD1-NTD and CTD.

**Fig. 3.**
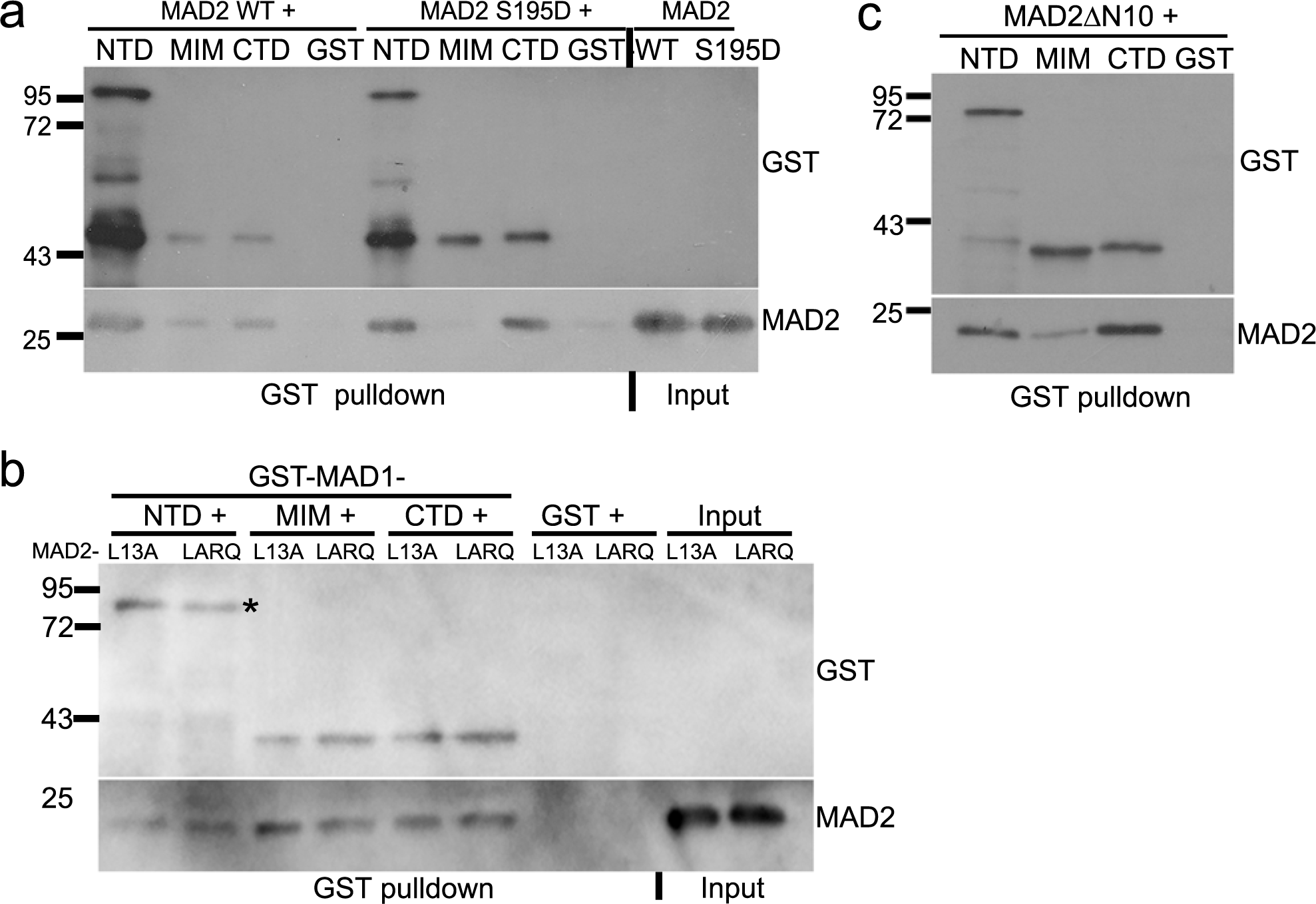
A novel interface in MAD2 is employed for its association with MAD1-NTD or –CTD. **(a)** GST-pulldown assays using GST-tagged MAD1 fragments after incubation with 6×His-tagged wild type MAD2 or MAD2-S195D mutant. **(b)** GST-pulldown assays using GST-tagged MAD1 fragments with MAD2^ΔN10^. **(c)** GST-pulldown assays using GST-tagged MAD1 fragments with MAD2^L13A^ and MAD2^LARQ^.

### MPS1 kinase phosphorylates MAD1-NTD and CTD

It is known that the kinase activity of MPS1 is essential for maintaining mitotic arrest even when MAD1 constitutively localizes at kinetochores as a fusion with mCherry-Mis12 (25–28). We hypothesize that MPS1 might phosphorylate MAD1 or MAD2 to regulate the efficiency of the MAD1:C-MAD2 catalyst. *In vitro* kinase assays found that MPS1 indeed phosphorylates MAD1-NTD and CTD (Fig. 4a). The specificity of the kinase assay was validated by that reversine, a previously characterized MPS1 inhibitor(44), reduces *in vitro* phosphorylation of an artificial MPS1 substrate myelin basic protein (Fig. 4a, compare lanes 1 and 2 in the right panel). MAD1-MIM and MAD2 conformers are not good substrates for MPS1 under the experimental condition. Interestingly, recombinant MPS1 kinase binds to MAD1-NTD or CTD but not MIM in the absence of ATP (Fig. 4b).

**Fig. 4.**
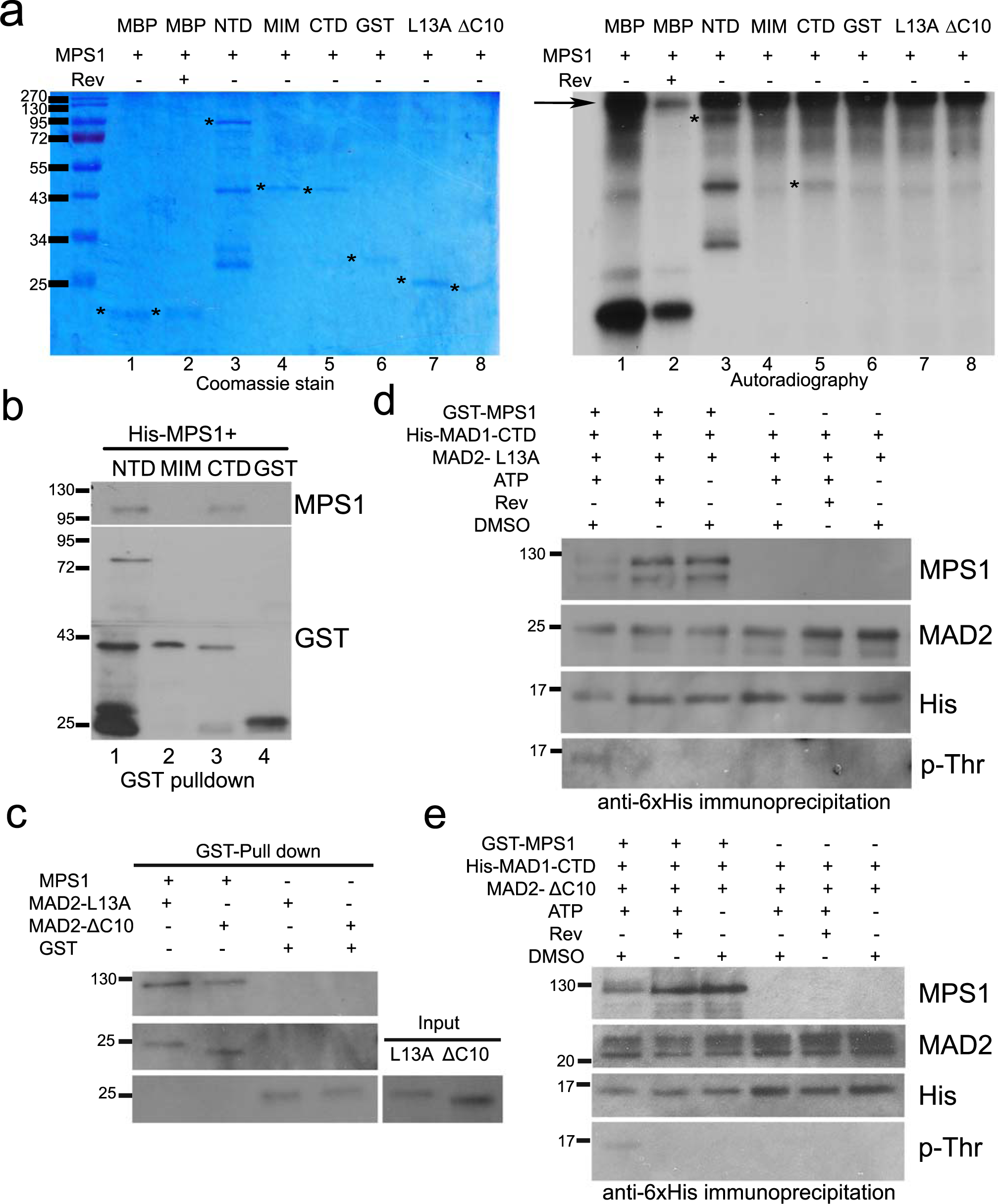
MPS1 phosphorylates MAD1 and interacts with MAD1 and MAD2. **(a)** *In vitro* kinase assays were performed using recombinant GST-MPS1, with myelin basic protein (MBP) as an artificial substrate of MPS1, and GST-MAD1-NTD, -MIM, -CTD, untagged MAD2^L13A^ or MAD2^∆C10^. The SDS-PAGE gel of the kinase assays were stained with Coomassie blue (left). Phosphorylation of the proteins by MPS1 was detected by autoradiography after the gel was dried (right). The asterisks indicate the expected sizes of corresponding proteins. The arrow indicates GST-MPS1. Reversine (Rev) was used in lane 2 to validate the kinase specificity. **(b)** GST-pulldown assays using GST or GST-tagged MAD1 fragments after incubation with His-tagged MPS1. **(c)** GST-pulldown assays using GST or GST-tagged MPS1 after incubation with His-tagged MAD2^∆C10^ or MAD2^L13A^. **(d)** and (**e**) Immunoprecipitation using anti-6×His antibody to detect His-tagged MAD1-CTD interaction with untagged MAD2^L13A^(d) or MAD2^∆d)^ (e) in the presence/absence of GST-MPS1 kinase, ATP or MPS1 inhibitor reversine.

We then tested whether MPS1 phosphorylation affects the interactions between MAD1-CTD and MAD2 conformers. We focused on CTD because it is not only easily purified without apparent degradation but also shows functional significance (see below). We did notice that GST-MPS1 could also bind to MAD2^L13A^ and MAD2^ΔC10^ (Fig 4c). *In vitro* incubation of MPS1, CTD and MAD2 led to phosphorylation of CTD, as evidenced by the mobility shift and positive signals of phospho-Thr antibody (lane 1 in Fig 4d and 4e). However, the association of either MAD2^L13A^ or MAD2^ΔC10^ with CTD did not show obvious changes, when compared to reactions in the absence of ATP or in the presence of MPS1 inhibitor reversine. The interactions between phosphorylated CTD and MPS1, nevertheless, became weaker (Fig. 4d and 4e).

### Functional characterization of MPS1 phosphorylation sites on MAD1

Realizing some limitations of *in vitro* binding experiments especially with protein fragments, we decided to directly test the potential effects of MPS1 phosphorylation sites on MAD1 in cells. Through mass spectrometry we identified 8 *in vitro* MPS1 phosphorylation sites on MAD1-NTD and CTD. The sites are T8, S22, S62, T323, S598, S610, T624 and T716 (Fig. 5a). To test the importance of these sites in cells, a mCherry-Mis12-MAD1^8A^ mutant with all the sites mutated into alanine was expressed and found to be defective in maintaining the mitotic arrest (229±23 min mitotic duration) (Fig. 5b). Mutating all four NTD sites into alanine showed no effect, but alanine mutants at the CTD four sites (MAD1^CTD4A^) shortened the mitotic duration comparable to MAD1^8A^ (Fig. 5b). Further tests found that little CTD^4A^ is phosphorylated when compared to wild type CTD (CTD^WT^) *in vitro*, supporting that the CTD four sites, S598, S610, T624 and T716, constituted primary phosphorylation sites of MPS1 kinase under our *in vitro* experimental conditions (Fig. S4a).

**Fig. 5.**
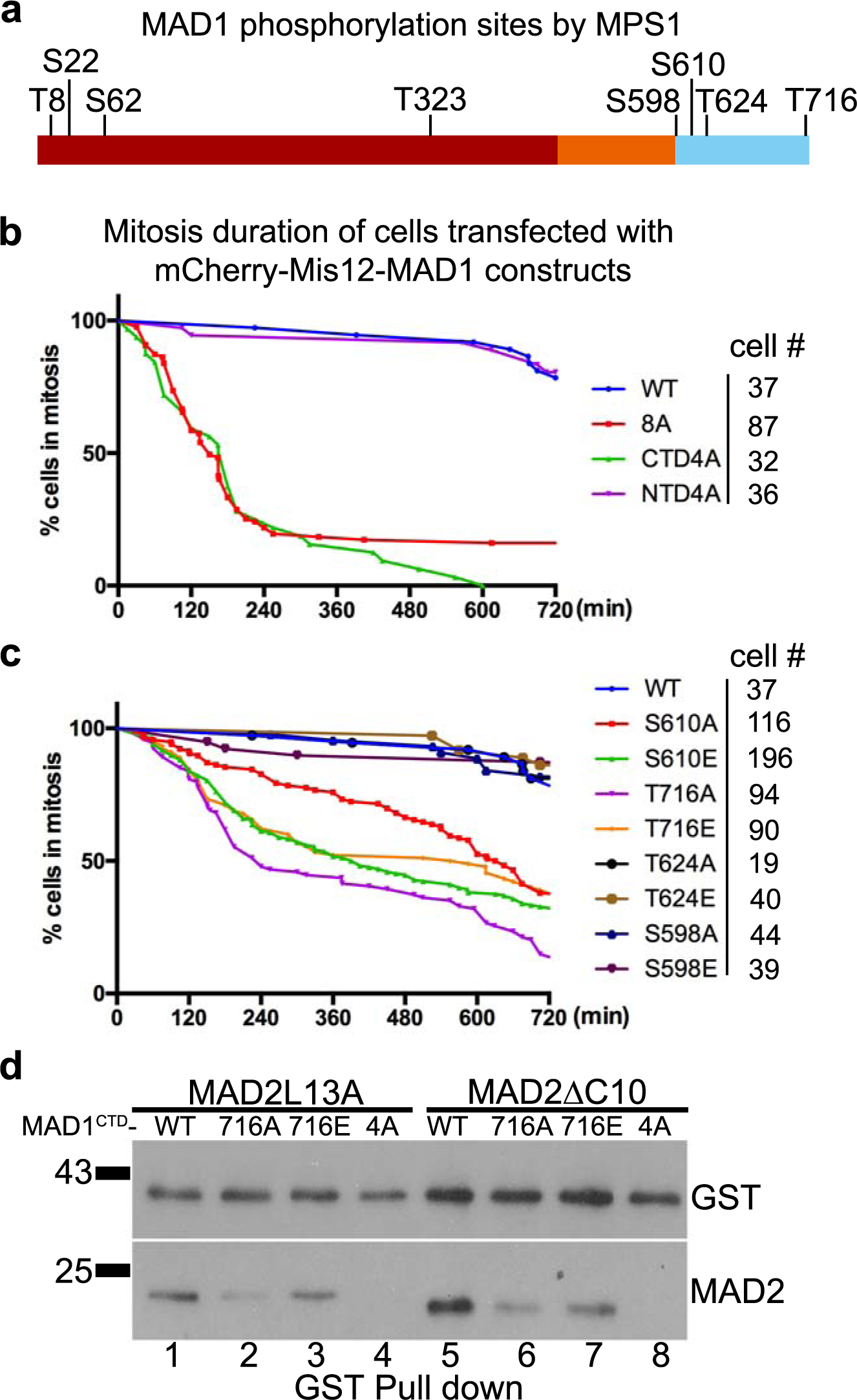
The putative MPS1 phosphorylation sites on MAD1-CTD are required for the mitotic checkpoint. **(a)** MAD1 phosphorylation sites by MPS1 kinase *in vitro* as determined by mass spectrometry. **(b)** Mitotic durations of HeLa cells transfected with mCherry-Mis12 fused with MAD1-WT, -8A (T8, S22, S62, T323, S598, S610, T624, T716 to A), -NTD4A (full length but T8, S22, S62, T323 to A) or -CTD 4A (full length but S598, S610, T624, T716 to A). Cell numbers imaged for each construct was listed on the right. **(c)** Phosphomimic and phospho-resistant mutants at individual MPS1 phosphorylation sites in the MAD1-CTD (S598, S610, T624, T716) were prepared as mCherry-Mis12 fusion constructs and transfected HeLa cells were imaged for mitotic durations as above. **(d)** GST-tagged MAD1-CTD fragments as wild type (WT), or containing T716A or T716E or CTD-4A mutations were incubated with either MAD2^L13A^ or MAD2^∆C10^, then GST pulldowns were probed with anti-GST and anti-MAD2 antibodies.

Next a mCherry-Mis12-MAD1^CTD4E^ phosphomimic mutant was prepared and surprisingly found to be also defective in maintaining the mitotic arrest (Fig. S4b). We reasoned that for a functional mitotic checkpoint in cells one or more of the four residues identified through *in vitro* kinase assays could not tolerate mutations. Both phospho-resistant and phosphomimic mutants at individual sites were then prepared. The mutants at S598 or S624 did not show obvious defects in the mitotic checkpoint when fused with mCherry-Mis12. However, the mutants at S610 and T716 showed significant differences in mitotic arrest durations as compared to the MAD1^WT^ fusion (Fig. 5c). Furthermore, the T716E mutant maintained the checkpoint longer than the T716A mutant (441±38 vs 242±48 min, P<0.05, students’ *t*-test), indicating that T716 is likely the residue activated by MPS1 for mitotic checkpoint signaling. However, the mCherry-Mis12 fusion of MAD1^T716E^ mutant could not maintain mitotic arrest when transfected cells were challenged by reversine (Fig. S4c), suggesting that either the phosphomimic mutant is imperfect or MPS1 has additional key substrates required for a functional mitotic checkpoint.

To further understand the imaging results with different CTD mutants, we examined whether these mutants affected the interaction of MAD1^CTD^ with MAD2. Both CTD^4A^ and CTD^4E^ showed defects in binding to O-MAD2 and C-MAD2 (Fig. S4d). Furthermore, CTD^T716A^ binds less O-MAD2 and C-MAD2 as compared to CTD^WT^ or CTD^T716E^ (Fig. 5d). Similarly, the functional defective MAD1^Y634E^ mutant also binds to less O-MAD2 and C-MAD2 than MAD1^Y634F^ (Fig. S4e). These results correlated mitotic arrest durations with the capability of MAD1^CTD^ mutants to bind to MAD2 (Fig. 1d, Fig. 5, and Fig. S4).

### MPS1 phosphorylation of MAD1 modulates inter-domain and inter-molecular interactions

Since MAD1-MIM, CTD and NTD are all required for an effective mitotic checkpoint (Fig. 1), we next investigated whether there is coordination between these domains. The presumed non-coiled coil domains along the length of MAD1 might provide certain bends and turns to the molecule, and an earlier model did suggested that MAD1^CTD^ folds back to the proximity of the catalytic core consisting of MIM and associated C-MAD2 (11). We examined and found that NTD directly interacts with CTD (Fig. 6a, lane 1). CTD interacts with itself (lane 3), agreeing with published crystal structure and acting as a positive control in this binding assay (33). MIM does not directly interact with CTD nor NTD (Fig. 6a, lane 2 and Fig. 6b). Addition of MPS1 kinase reduced the interactions between MAD1^NTD^ and MAD1^CTD^ (Fig. 6c). The effect depends on the MPS1 kinase activity as omitting ATP from the reactions or adding MPS1 inhibitors (reversine or AZ3146) reversed the MPS1 effect on NTD:CTD interaction (compare His signals in lane 2 and those in lanes 3-5).

**Fig. 6.**
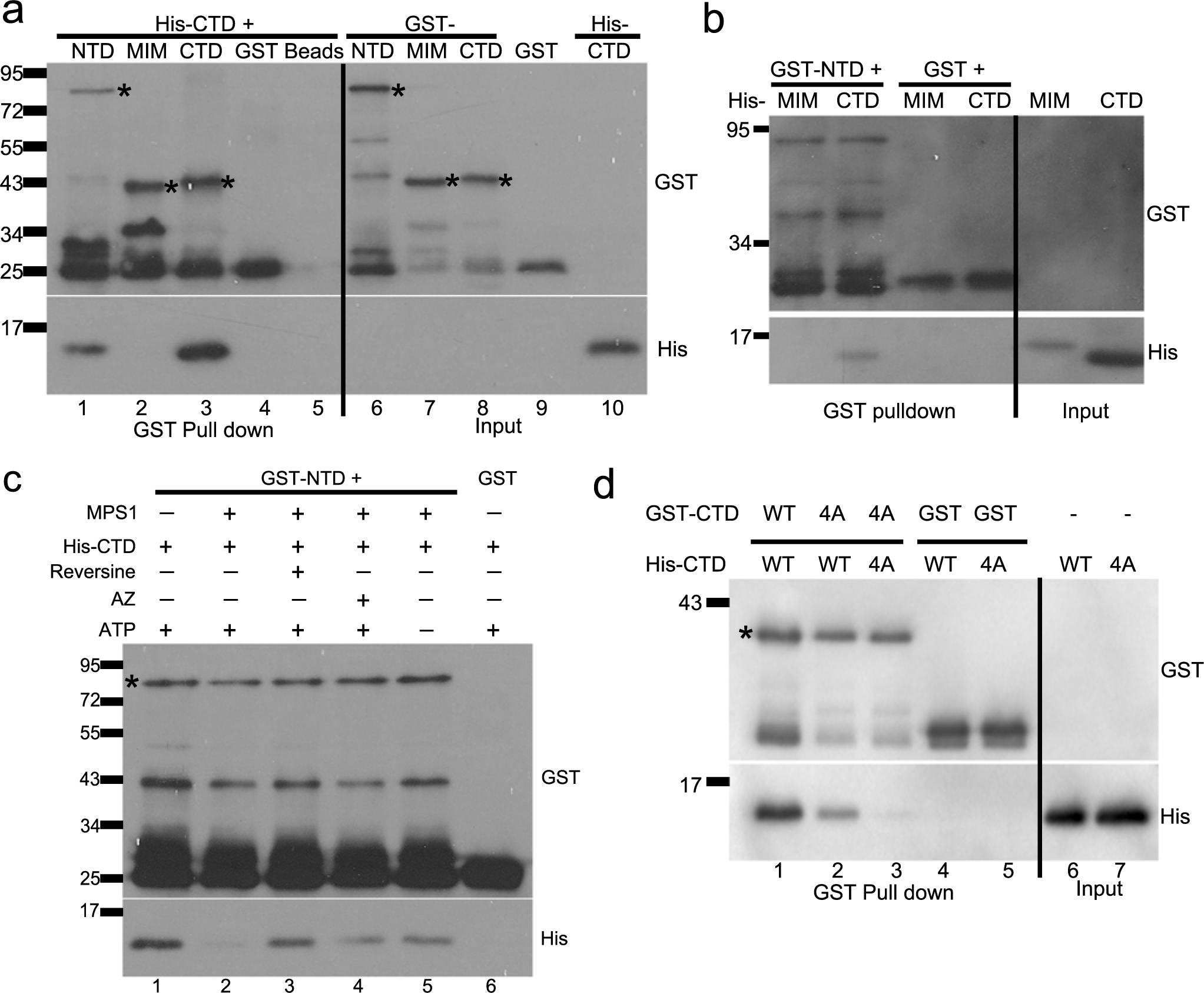
Inter-domain interactions of MAD1 are modulated by MPS1 kinase. **(a)** GST-tagged MAD1-NTD, -MIM, -CTD, GST or beads alone were incubated with 4µM of His-tagged MAD1-CTD. The GST pulldown samples were separated by SDS-PAGE and subjected to Western blot using anti-GST and anti-6×His antibodies. The asterisks marked the expected bands of GST-MAD1-NTD,-MIM and -CTD. **(b)** GST pulldown assays after GST or GST-MAD1-NTD was incubated with His-tagged MAD1-MIM and MAD1-CTD. **(c)** Recombinant GST-MAD1-NTD or GST were incubated with 9 µM of His-MAD1-CTD in the absence or presence of MPS1 kinase, reversine (MPS1 inhibitor), AZ3146 (another MPS1 inhibitor) and ATP. GST pulldowns were then probed for anti-GST and anti-6×His antibodies. **(d)** GST-MAD1-CTD^WT^ or -CTD^4A^ was incubated with His-tagged MAD1-CTD^WT^ or -CTD^4A^ and GST pulldowns were probed.

MAD1 dimerization has been observed in MAD1^MIM^:MAD2 and MAD1-CTD crystal structures (11,33). However, whether MAD1 dimerization is essential for its activity to promote MAD2 O-C conversion is unclear. We noticed that in Fig. 1e and 1f, MAD1 truncations missing either NTD or CTD did not co-immunoprecipitate endogenous MAD1 as well as other fusions. Furthermore, recombinant GST-CTD^4A^ fragment also binds less His-CTD^WT^ and even less His-CTD^4A^ (Fig. 6d). These results suggested a possible role of CTD in MAD1 dimerization or oligomerization (see discussion).

## DISCUSSION

MAD2 O-C conversion is a key signal amplification step for the mitotic checkpoint. Current model suggests that the conversion is catalyzed by a unusual catalyst: the MAD1:C-MAD2 complex localized at unattached kinetochores. How the catalysis is achieved is still unclear. Together with two very recent publications (29,30), our results support the roles of MAD1-CTD and MPS1 kinase in promoting the MAD2 O-C conversion. Our work agreed with the critical role of MAD1^T716^ which is phosphorylated by MPS1, but also suggested that S610 and Y634 are potential key residues for regulating MAD2 O-C conversion (Fig 1, Fig 5). S610 is phosphorylated by MPS1 *in vitro* (Fig 4), and Y634 was reported to be phosphorylated *in vivo* although we have not confirmed this yet in mitotic cells (37). We have also uncovered additional protein-protein interactions between MPS1, MAD1 and MAD2 (Figs. 2, 3, 4, 6). Of particular interest, we found that MAD1-NTD and CTD interact with each other and both bind to O-MAD2 and C-MAD2. These results have been integrated into an updated model for the MAD1:C-MAD2 catalyst that promotes MAD2 O-C conversion (Fig. 7). Many mechanistic details remain to be filled, so we hereby discuss the implications of our results.

**Fig. 7.**
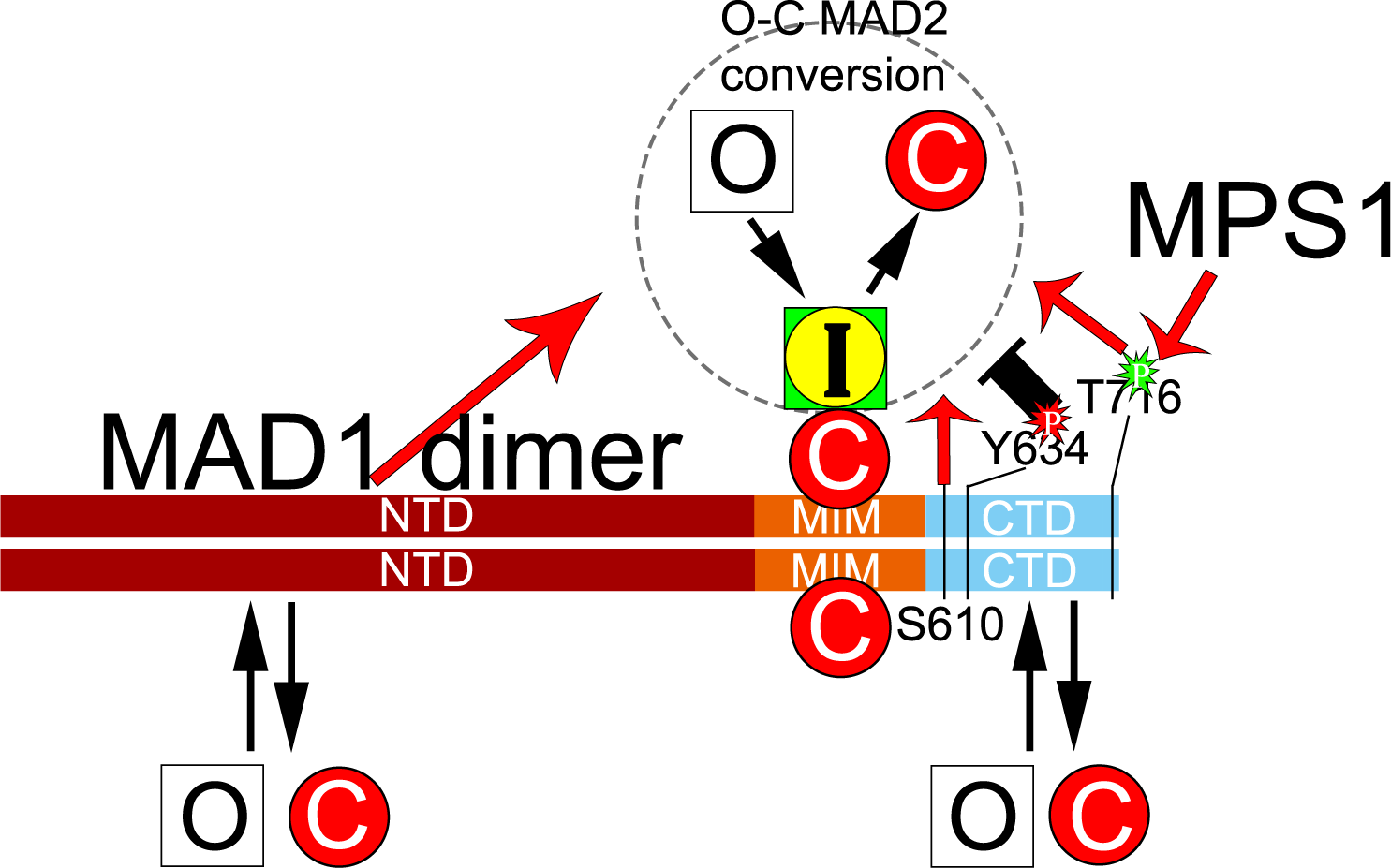
An updated model on the MAD1:C-MAD2 catalyst for MAD2 O-C conversion. In the classical model, a MAD1 dimer tightly binds two C-MAD2 molecules at its MIM region to become the catalyst for MAD2 O-C conversion. Our work showed that MAD1-NTD and CTD have additional, probably weaker, binding sites for both O-MAD2 and C-MAD2. Cell imaging results suggested that MAD1-NTD and CTD positively contribute to the MAD2 O-C conversion. Furthermore, the possible functions of several residues in the CTD domain were revealed. T716 phosphorylation by MPS1 and S610 are required for full MAD1 activity. Y634 might be a residue whose phosphorylation negatively impacts the O-C conversion. Similarly, the NTD:CTD interaction (not shown) may restrain the catalytic efficiency of MAD1, but the interaction can be disrupted by action of MPS1 kinase. It remains unclear whether both C-MAD2 molecules bound to the MIM regions are equally engaged in MAD2 O-C conversion.

### The functions of MAD1-NTD and CTD in MAD2 O-C conversion

The crystal structure of MAD1-MIM in complex with C-MAD2 has solidified the now classical model of the MAD1:C-MAD2 catalyst as a 2:2 heterotetramer (11) (Fig S1). Each of the two “liganded” C-MAD2 tightly wraps around one MIM monomer through its “safety belt” loop (11,18,38). C-MAD2 then utilizes its dimerization domain to recruit O-MAD2 and converts the latter into C-MAD2, resulting in signal amplification for the mitotic checkpoint (6,7,12). The conversion may go through multiple intermediate states (I-MAD2) (6,7,10,13,32). At least two structures of I-MAD2:C-MAD2 dimers have been solved, one possibly representing the initial O-MAD2 “docking” complex and the other containing a later stage of I-MAD2 state (MAD2^ΔN10^) (10,32). The major structural features distinguishing O- and C-MAD2 lie at the N- and C-termini (8). The solved MAD2^ΔN10^ structure indicates a topology similar to O-MAD2 but its internal core more closely resembles that of C-MAD2 (10). However, one long suspected intermediate state with partially unfolded N- or C-termini, which is more amenable for MAD2 refolding or its interaction with CDC20, has not been identified based on studies of various MAD2 conformational dimers therefore was considered as only a fleeting state (10). Such a state, as suggested (29,30), may be coupled with formation of the CDC20:MAD2 complex or the MCC, making the MAD2 O-C conversion reaction more rapid and energy favorable. The coupling could be achieved at unattached kinetochores when either BUB1 or T716 phosphorylated MAD1 binds and presents CDC20 to I-MAD2 produced by the MAD1-MIM:C-MAD2 complex (29,30).

Although much of MAD1 has been usually simplified as rather stiff coiled coils, the known structures of MAD1^MIM^ and MAD1^CTD^ and careful analyses of MAD1^NTD^ showed that multiple MAD1 segments may adopt alternative structures or remain disordered (11,33) (Fig. 1, Fig. S1). It is likely that “liganded” C-MAD2 binding to the MIM region of MAD1 still constitutes the catalytic core and represents the stably MAD1-associated MAD2 in cells (Fig. 1e), but earlier work by others have already lent strong support to possible MAD1-CTD involvement in the mitotic checkpoint responses (12–18,29,30). At least part of the MAD1-NTD (1-485 residues) contributes to MAD1 nuclear pore or kinetochore localization and interaction with other proteins including Ndc80, Plk1, Nek2A, Tpr, Cep57 and CENP-E (20–23,33,45–49). The regions spanning (400-500) or (420-485) could possibly impact on the mitotic checkpoint (14,29). Nevertheless, the possibility that MAD1-NTD and -CTD directly impacts on MAD2 O-C conversion was not thoroughly studied until this work.

Our results suggest that a full length MAD1 molecule may employ NTD and CTD to at least transiently interact with the substrates (O-MAD2) and the products (C-MAD2) even the intermediate states (I-MAD2, represented by MAD2^ΔN10^ in Fig. 3C) of the MAD2 O-C conversion reaction(50). The interaction with O-MAD2 might increase local concentration of substrates, driving the conversion reaction. Similarly, enrichment of I-MAD2 by MAD1-NTD or CTD could facilitate its interaction with CDC20 (29,30), although conversion to C-MAD2 can certainly be accomplished in the absence of CDC20 (12,13). Following the law of mass action, the NTD or CTD-retained C-MAD2 could also drive its interaction with BUBR1 as a step of MCC assembly (36). These possible scenarios may underlie the compromised mitotic checkpoint responses as seen when cells were transfected with mCherry-Mis12-MAD1^ΔNTD^ or - MAD1^ΔCTD^ (Fig. 1). In consistence with this idea, our data indicated that MAD1-NTD and MAD1-CTD bind to a new interface on MAD2. Such a binding mode is postulated to be advantageous as the I-MAD2 or C-MAD2 products anchored on MAD1 then have two other protein-protein interaction interfaces readily available: the safety belt for CDC20 and the dimerization domain for BUBR1 to assemble into the MCC, the anaphase onset inhibitor (1,4). In addition, the MAD2 interactions with NTD or CTD must be weak or transient in cells as mCherry-Mis12-MAD1^ΔNTD^ and MAD1^ΔCTD^ did not show significant reduction in co-immunoprecipitated MAD2 and FLAG-MAD1^NTD^ only shows faint MAD2 binding (Fig. 1). Such transient interactions are likely beneficial for stepwise relay mechanisms to facilitate MAD2 O-C conversion.

In the MAD1^MIM^:C-MAD2 tetramer structure (11,32), the dimerization interfaces along the two C-MAD2 molecules face towards opposite directions to recruit O-MAD2. It is therefore possible that endogenous full-length MAD1, when bound by C-MAD2 at MIM to form a catalyst, contains more than one catalytic center. It remains to be seen whether MAD1-NTD or CTD asymmetrically affects the two presumable MAD1^MIM^:C-MAD2 catalytic centers or directly contribute to the catalysis such as stabilizing the I-MAD2 with transiently unfolded N- or C-termini. In addition, MAD1-CTD and MAD1-NTD are important for MAD1 oligomerization (Fig. 1 and Fig. 6d). We noticed an earlier publication suggesting that S214 phosphorylation by ATM affects MAD1 dimerization (51). Whether MAD1 dimerization regulates MAD2 conversion is interesting to further explore in the future.

### The impact of mitotic kinases on MAD2 O-C conversion

Whatever the catalytic mechanism the MAD1:C-MAD2 complexes employ to promote MAD2 O-C conversion, the catalysis is likely initiated or enhanced by mitotic checkpoint kinases whose activities have been shown important for the checkpoint, including MPS1, BUB1 and Aurora B (25,26,28,31). These kinases not only help enrich the MAD1:C-MAD2 catalysts to unattached kinetochores, but also directly impact their catalytic efficiency. In this work we have focused on potential effect of MPS1 on increasing the catalytic efficiency of the MAD1:C-MAD2 complex. We expected to see differential bindings of O-MAD2 and C-MAD2 to MAD1-NTD or CTD, especially after MPS1 phosphorylation, but did not have experimental evidence to support the idea yet (Fig. 4). We wonder whether using full-length MAD1 or including additional kinases might generate different results (29,30), however, current work still provides interesting points for discussion. MPS1 is a key upstream kinase orchestrating the organization of other mitotic checkpoint proteins at unattached kinetochores. It phosphorylates KNL1 to recruit BUB1:BUB3 and BUBR1:BUB3 complexes (reviewed in (1,52)). It also phosphorylates BUB1 to recruit MAD1:MAD2 (30,31,53). Furthermore, MPS1 phosphorylating MAD1 at T716 may enhance MAD1 binding to CDC20 (30). Thus MPS1 activity may promote the formation of the MCC by placing all its components BUBR1:BUB3, CDC20 and MAD2 in spatial proximity (29,30). CDC20 binding to BUB1 or BUBR1 might also help the MCC assembly at unattached kinetochores (54–56).

We have demonstrated that MPS1 interacts with and phosphorylates MAD1-NTD and CTD (Fig. 4). Phospho-resistant mutants at *in vitro* MPS1 phosphorylation sites in MAD1-CTD especially the T716A mutation compromised the mitotic checkpoint responses in cells and showed reduced interaction with MAD2 *in vitro* (Fig 5). No increase in MAD2 interactions with MAD1-CTD was detected either using phospho-mimic T716E mutant or after direct *in vitro* phosphorylation by MPS1 (Fig. 4, Fig. 5). We cannot exclude the possibility that in cells and in the context of full length MAD1, CTD phosphorylation show differential binding towards either conformer of MAD2. Phosphorylation by MPS1 did reduce its own interaction with CTD and the interaction between NTD and CTD (Fig. 4, Fig. 6). We propose that the NTD:CTD interaction occurs in interphase cells and represents an inactive state of MAD1 even though its MIM associates with C-MAD2 in a cell cycle independent manner (18,24). The interaction between MPS1 and MAD1 might contribute to the MAD1 recruitment to kinetochores. Once concentrated at unattached kinetochores, MPS1 kinase becomes activated and phosphorylates BUB1 to stably anchor MAD1 (30,31,57). Activated MPS1 also phosphorylates MAD1-CTD, most likely at T716, relieving itself and the NTD from CTD. The now exposed and phosphorylated CTD could bind to O-MAD2 and C-MAD2 and also facilitate MAD2 O-C conversion through increased association with CDC20 (30) or other more direct mechanisms as discussed in last section.

Although deleting MAD1-NTD compromised mitotic checkpoint when examined using the mCherry-Mis12 fusions (Fig. 1), mutating presumable MPS1 phosphorylation sites within MAD1-NTD had no impact on the mitotic checkpoint (Fig. 5), and others found no requirement for MAD1 N-terminal 400 or 420 residues for MAD2 O-C conversion or checkpoint responses (14,29). This suggests that a functional critical region lies within (420-485) residues, which happens to contain a likely non-coiled coil structure (Fig. 1). However, neither of the MAD1 fragments spanning (327-423) or (327-488) residues bind to MAD2 *in vitro* (Fig. 2), indicating the positive effect of MAD1-NTD on the mitotic checkpoint signaling needs further clarification.

Interestingly Faesen et al reported that MAD2 can be phosphorylated by MPS1 *in vitro* at S195 (29). Previously phosphorylation at this site was proposed to render MAD2 hard to convert thus to more likely stay in O-conformation (43,58). Whether this particular MAD2 phosphorylation by MPS1 represents a feedback regulatory mechanism can be further explored. In this regard, it might be useful to note that MPS1 interacts with C-MAD2 and O-MAD2 while MAD2^S195D^ binds to MAD1 at NTD or CTD but not MIM (Fig 3, Fig 4).

Although recent work has proposed how BUB1 might help the MAD2 O-C conversion and MCC assembly (29–31), it will still be interesting to further define how other kinases such as Aurora B regulate the MAD1:C-MAD2 catalyst to increase C-MAD2 production when the checkpoint becomes activated in cells. In addition, there have been reports that some tyrosine kinases may help silence the mitotic checkpoint to drive anaphase onset (59–63). MAD1 Y634 seems to play important roles in the mitotic checkpoint (Fig 1, Fig S4) and it was once reported to be phosphorylated *in vivo* (37). Further characterization of this site as well as the apparently essential S610 and T716 residues at the MAD1-CTD will provide more mechanistic insights into the functions of MAD1 in MAD2 O-C conversion.

## EXPERIMENTAL PROCEDURES

### DNA Constructs

The MAD1 and MAD2 DNA fragments and mutants used in the work are summarized in Table S1. The pMSCV-mCherry-Mis12-MAD1 constructs with wild-type or a mutant MAD1 (K541A, L543A) were gifts from Dr. Tarun Kapoor (Rockefeller University) (26,28). Other MAD1 mutants or truncations were cloned or mutated in an intermediate pENTR2B vector (Invitrogen), cut out as a *Not*I-*EcoR*I fragment, and then used to replace the *MAD1* gene in the pMSCV-mCherry-Mis12 backbone also cut with *Not*I and *EcoR*I. Mutagenesis was conducted using QuickChange Lightning Multi Site-Directed Mutagenesis Kit (Agilent). Different fragments of MAD1 were also cloned into a pENTR-D-TOPO vector using TOPO cloning kit (Invitrogen) and then recombined into pDEST15 (for N-terminal GST tag) or pDEST17 (for N-terminal 6×His tag) vectors through LR reactions as instructed for the Gateway cloning system (Invitrogen). MAD2 expression constructs were prepared similarly as before (36), but with a TEV cleavage site inserted between the tags and second MAD2 codon during pENTR cloning steps. MPS1 was recombined into pDEST10 or pDEST20 and transformed into DH10BAC to prepare recombinant bacmids for Sf9 insect cell transfection. All constructs were confirmed by DNA sequencing.

### Cell Culture and Transfection

HeLaM, a subline of HeLa (35), was maintained in DMEM with 10% fetal bovine serum at 37°C in 5% CO_2_. DNA transfection was carried out using TransIT-LT1 reagent (Mirus) following the manufacturer’s instructions or using polyethylenimine (PEI) as described before (64). Sf9 cells were grown at 27 °C in SFX medium (Hyclone) in the presence of antibiotics (streptomycin/penicillin). Cellfectin (Invitrogen) was used to transfect bacmids into Sf9 cells.

### Live Cell Imaging

For determining mitotic durations, HeLa cells grown on No. 1.5 coverslip-bottomed 35mm dishes (MatTek) were transfected with different mCherry-Mis12-MAD1 constructs. Usually 24 hr after transfection, live cell imaging was started on an automated Olympus IX-81 microscope to collect phase contrast and RFP images at 15 min intervals using a 60 × objective lens (NA=1.42) while cells were maintained at 37 °C in a heating chamber. Single-plane images were acquired for up to 13 hr at multiple positions using a CoolSNAP HQ2 camera with 2×2 binning. Student’s *t*-test was used to evaluate the statistical significance between the differences in mitotic durations after different treatment. Nuclear envelope breakdown marks the beginning and appearance of cleavage furrow the end of a mitosis. Some images were collected on a Leica TCS SP8 confocal microscope with a 63 × objective (NA = 1.40) as *z*-stacks of 1.0 μm.

### Cell lysates, immunoblotting and immunoprecipitation

These were performed as described before (28,64). A list of primary antibodies used in this study is summarized in Table S2.

### Recombinant proteins

GST-tagged or His-tagged MAD1 fragments and His-TEV-tagged MAD2^L13A^ or His-TEV-tagged MAD2^∆C10^ were expressed in *E. coli* BL21(DE3)-CodonPlus RIPL (Stratagene), normally at 16°C. His-MPS1 or GST-MPS1 was expressed in Sf9 cells after infection with recombinant baculoviruses. All expressed proteins were purified using GSH-agarose or Probond nickel beads (Invitrogen). His-tagged TEV(S219P) protease(65) was used to cleave His-tag to make untagged MAD2. Peak fractions of eluted proteins were pooled, buffer-exchanged and concentrated using Pierce protein concentrators with 10k cutoff. The storage buffer is 50mM Hepes pH7.5, 100mM KCl, 1mM DTT, 30% glycerol. Concentrations of recombinant proteins were determined by comparing the target band with BSA standards on Coomassie blue stained gels using ImageJ software (66).

### *In vitro* binding assays

4µl of 5× binding buffer (100mM Tris-HCl, pH 8.0, 750mM NaCl, 50mM MgCl_2_, 2.5% NP-40, 50% glycerol, 500 µg/ml BSA) was mixed with recombinant GST-tagged MAD1 fragments and MAD2 mutants or His-MAD1-CTD or His-MPS1. The proteins were used at roughly endogenous intracellular concentrations unless stated otherwise in the figure legends: [MAD2]=230 nM, [MPS1]=100 nM, and [MAD1]=60 nM. H_2_O was added to make the final volume of 20µl. The reactions were incubated at 37 °C for 1 hr and then mixed with 10µl GSH agarose beads and rotated at 4°C for 40 min. The beads were pelleted and washed 4 times with wash buffer (1× PBS, pH 7.4, 150 mM NaCl, 0.5% NP-40, 10% glycerol). Then 10µl 2× SDS sample buffer was added to the beads and the samples were heated at 80°C for 10 min before SDS-PAGE. The gel was transferred to PVDF membranes (Millipore) for immunoblotting.

### *In vitro* kinase assays

GST-tagged MPS1 kinase was purchased from Invitrogen or was purified from Sf9 cell lysates as well as His-tagged MPS1 (64). Myelin basic protein (MBP) was purchased from Sigma as an artificial substrate for MPS1. *In vitro* kinase assays were set up similarly as previously described (28,64): 4 µl of 5× kinase buffer (125 mM Tris-HCl, pH 7.5, 300 mM β-glycerophosphate, 50 mM MgCl_2_) was mixed with recombinant kinase, substrates, 5µCi ^32^P –ATP and 50 µM cold ATP. In some reactions, MPS1 kinase inhibitors reversine (Calbiochem) or AZ3146 (Selleckchem) was used at 500 nM and 2 µM final concentrations, respectively. DMSO concentration is kept below 0.5%. H_2_O was added to make the final volume of 20 µl. The reactions were incubated at 30°C for 30 min and then terminated by adding 20 µl 2×SDS sample buffer. Samples were subjected to SDS-PAGE followed by Coomassie staining. After destaining, the SDS-PAGE gel was vacuum dried. Phosphorylation of the substrates was visualized by autoradiography. Samples for mass spectrometry were prepared using only 0.5 mM cold ATP in the kinase reactions and separated on SDS-PAGE. Phosphorylated residues on the excised bands were determined by mass spectrometry (MSBioworks, Ann Arbor).

## ACKNOWLEDGMENTS

We thank Drs. Tarun Kapoor and Maria Maldonado for the mCherry-Mis12-MAD1-WT and -AA constructs and Dr. Tim Yen for multiple antibodies. We also thank Dr. William Taylor, Dr. Qian Chen and the members of the Liu lab for stimulating discussions. STL is funded by NIH R01CA169500.

## CONFLICT OF INTEREST

The authors declare no conflict of interest.

## AUTHOR CONTRIBUTIONS

S.T.L. conceived and supervised the project. S.T.L. and W.J. designed the experiments with input from Y.L. and E.A. W.J. did most of the experiments. Y.L. performed cloning and mutagenesis of some MAD1 mutants, participated in protein purification, *in vitro* binding and *in vitro* kinase assays and live cell imaging. E.A. carried out some protein purification and *in vitro* binding experiments. S.T.L. drafted the manuscript and all authors contributed to finalizing the manuscript.

## FOOTNOTES

1 The abbreviations used are: O-MAD2: open MAD2; C-MAD2: closed MAD2; NTD: N-terminal domain; CTD: C-terminal domain; MCC: mitotic checkpoint complex; APC/C: anaphase promoting complex/cyclosome; MIM: MAD2 interaction motif; TEV: tobacco etch virus; SDS-PAGE: SDS polyacrylamide gel electrophoresis; MBP: myelin basic protein; WT: wild type.

